# Structure of the herpes-simplex virus portal-vertex

**DOI:** 10.1101/290544

**Authors:** Marion McElwee, Swetha Vijayakrishnan, Frazer Rixon, David Bhella

**Affiliations:** Medical Research Council – University of Glasgow Centre for Virus Research, Sir Michael Stoker Building, Garscube campus, 464 Bearsden Road, Glasgow G61 1QH, United Kingdom

**Keywords:** herpesvirus, capsid, virus structure, cryoEM, asymmetry, portal, DNA, genome

## Abstract

Herpesviruses include many important human pathogens such as herpes simplex virus, cytomegalovirus, varicella-zoster virus and the oncogenic Epstein-Barr virus and Kaposi-sarcoma associated herpesvirus. Herpes virions contains large icosahedral capsids that have a portal at a unique five-fold vertex, similar to that seen in the tailed bacteriophages. The portal is a molecular motor through which the viral genome enters the capsid during virion morphogenesis. The genome also exits the capsid through the portal-vertex when it is injected through the nuclear-pore into the nucleus of a new host cell to initiate infection. Structural investigations of the herpesvirus portal-vertex have proven challenging, owing to the small size of the tail-like portal-vertex associated tegument (PVAT), and the dense tegument layer that lays between the nucleocapsid and the viral envelope, obscuring the view of the portal-vertex. Here we show the structure of the herpes simplex virus portal-vertex at sub-nanometer resolution, solved by electron cryomicroscopy (cryoEM) and single-particle 3D reconstruction. This led to a number of new discoveries including the presence of two previously unknown portal associated structures that occupy the sites normally taken by the penton and the Ta triplex. Our data revealed that the PVAT is composed of ten copies of the C-terminal domain of pUL25, which are uniquely arranged as two tiers of star-shaped density. Our 3D reconstruction of the portal-vertex also shows that one end of the viral genome extends outside the portal in the manner described for some bacteriophage but not previously seen in any eukaryote viruses. Finally, we show that the viral genome is consistently packed in a highly-ordered left-handed spool to form concentric shells of DNA. Our data provide new insights into the structure of a molecular machine critical to the biology of an important class of human pathogens.

## Introduction

Herpes Simplex Virus 1 and 2 (HSV-1 and HSV-2) are important human pathogens. It is estimated that ∼90% of the world’s population are infected with one or both viruses^1^. HSV-1 is the primary cause of cold sores, and HSV-2 of genital herpes. These conditions are both highly contagious, and HSV-2 is amongst the most common sexually transmitted infections. Infection with HSV is lifelong, owing to the ability of herpesviruses to enter a latent state with periodic reactivations^2^. HSV can also cause more serious conditions including keratitis, which may lead to loss of sight^3^, and a potentially fatal encephalitis^4^. The herpesvirus family includes many other important human pathogens, such as varicella-zoster virus the cause of chicken pox and shingles; cytomegalovirus a notable cause of congenital abnormalities; Kaposi-sarcoma associated herpesvirus that causes cancer in immune-compromised individuals and Epstein-Barr virus, the cause of infectious mononucleosis that has also been linked to several cancers.

Herpesviruses are large double stranded DNA viruses, having genomes up to 240 kbp. The viral DNA is packaged in a complex T=16 icosahedral capsid that is 1250Å in diameter^5^. The DNA containing capsid, or nucleocapsid, is embedded in a dense proteinaceous layer known as the tegument that is in-turn surrounded by a host-derived lipid envelope. The viral envelope is studded with glycoproteins that mediate viral attachment and entry. HSV virions enter host cells by fusing their envelopes with the host cell plasma membrane, allowing the nucleocapsid and tegument to enter the cytoplasm^6^. The nucleocapsid traffics along microtubules to the microtubule organising centre, and from there, to the nucleus^7^. The nucleocapsid then docks to a nuclear-pore complex, through which it injects its genome into the nucleus^8,9^. DNA egress from the capsid is through a unique portal-vertex, located at an icosahedral five-fold symmetry axis.

The portal-vertex is also the means by which the viral DNA is packaged into nascent capsids within the nucleus^10^. Virion morphogenesis commences in the nucleus with the formation of the procapsid; an icosahedrally symmetric spherical shell assembly of capsomeres that are hexamers (hexons) and pentamers (pentons) of the major capsid protein pUL19 (VP5)^11^. Heterotrimers of pUL38 (VP19C) and pUL18 (VP23) – termed triplexes, along with the scaffold protein pUL26 direct procapsid assembly, which nucleates around the dodecameric portal formed by pUL6^12^. Procapsid maturation: angularisation and expulsion of the scaffold protein occurs as the viral genome is pumped into the shell^13^. Replication of the viral genome results in formation of a concatemer, from which unit-length genomes are packaged into procapsids by the portal (pUL6) and terminase complex consisting of pUL33, pUL28 and pUL15^14,15^. Cleavage of the DNA, by the terminase, occurs upon recognition of specific signals. The capsid associated tegument complex (CATC – previously termed CCSC and CVSC), comprised of pUL17, pUL25 and pUL36 binds to the triplexes and hexons about the icosahedral five-fold vertices of mature capsids within the nucleus^16-21^. Notably, it has been observed that pUL25 is essential for retention of DNA within the nucleocapsid^22,23^. Moreover, pUL25 has also been shown to be important for genome release^24^.

The mature nucleocapsid leaves the nucleus by budding through the nuclear membrane via an envelopment/de-envelopment step^25^. Cytoplasmic capsids acquire further tegument proteins in the cytoplasm and are enveloped by budding into plasma membrane derived lipid vesicles, from which they are released by exocytosis at the cell surface^26^.

The structure of the HSV particle has been the subject of investigation for over thirty years^27^, using cryoEM and icosahedral 3D reconstruction to determine the high-resolution features of the nucleocapsid^5,21,28^, and lower resolution tomography to investigate the nature of asymmetric features such as the portal-vertex and viral envelope^29,30^. Attempts to resolve the structure of the portal-vertex have been largely unsuccessful however, owing to the dense tegument layer that obscures this feature, combined with the high symmetry of the viral capsid, which dominates attempts to align particle images for asymmetric reconstruction.

Here, we show the structure of the portal-vertex of HSV-1 at 8 angstroms resolution, revealed by focussed-classification and 3D reconstruction of cryoEM images of purified virions. These data reveal that the usual pUL19 penton is replaced by a unique five-fold symmetric assembly. This feature displays five well-defined coiled-coil motifs, each made up of two α-helices, arranged perpendicular to the capsid surface about the five-fold symmetry axis. It appears to be anchored to the virion by interactions with triplex-like structures that occupy the position normally taken up by peri-pentonal Ta triplexes, immediately about the five-fold vertex. The CATC assembly is still present and, similarly to penton associated CATC, is bound to the Tc triplexes, forming a bridge across the peri-portal triplex-like structures towards the five-fold axis. We interpret our data as showing that the pUL25 C-terminal domains are positioned differently to those seen at penton-vertices, giving rise to a small tail-like assembly that crowns the unique five-fold vertex. Strong density was seen to extend through the portal-vertex structures that we interpret as DNA. This suggests that the trailing end of the packaged genome remains engaged in the portal-vertex, ready for release through the nuclear-pore. Our reconstruction also reveals the arrangement of packaged DNA within the virion, which is clearly resolved as a left-handed spool arranged in concentric layers.

We provide the highest-resolution view to date of a critical component of the herpesvirus virion. The portal is a molecular machine responsible for both packaging and release of the viral genome, in one of the most important groups of viral pathogens to infect humans. Furthermore, our focussed classification approach demonstrates the power of modern image processing algorithms, to break the shackles of symmetry that have limited our understanding of virus structural biology for so long.

## Results

To determine the structure of the unique five-fold vertex comprising the portal and portal-vertex associated tegument (PVAT), we imaged purified HSV-1 virions by cryo-electron microscopy (figure S1). A total of 3,702 micrographs were captured from which a dataset of 6,069 virion images was extracted for 3D reconstruction. An initial 3D reconstruction was calculated with full icosahedral symmetry imposed, in order to accurately define the particle origins and orientations in each image (figure 1a). The icosahedral reconstruction achieved a resolution of 6.3 Å (figure S2) and closely resembled recently published structures at a similar resolution, revealing well defined CATC density with a clear four-helix bundle that has been attributed to pUL17, pUL25 and pUL36 (figure 1b,f)^28^. At lower isosurface threshold, we see two distinct globular domains per CATC, one on top of the pUL19, and one lying to the side. These regions have been convincingly assigned as C-terminal domains of pUL25 in the closely related pseudorabies virus^16^, which displays similar density to the CATC of HSV-1, thus there are ten pUL25 molecules per five-fold vertex (figure 1e).

**Figure 1.**
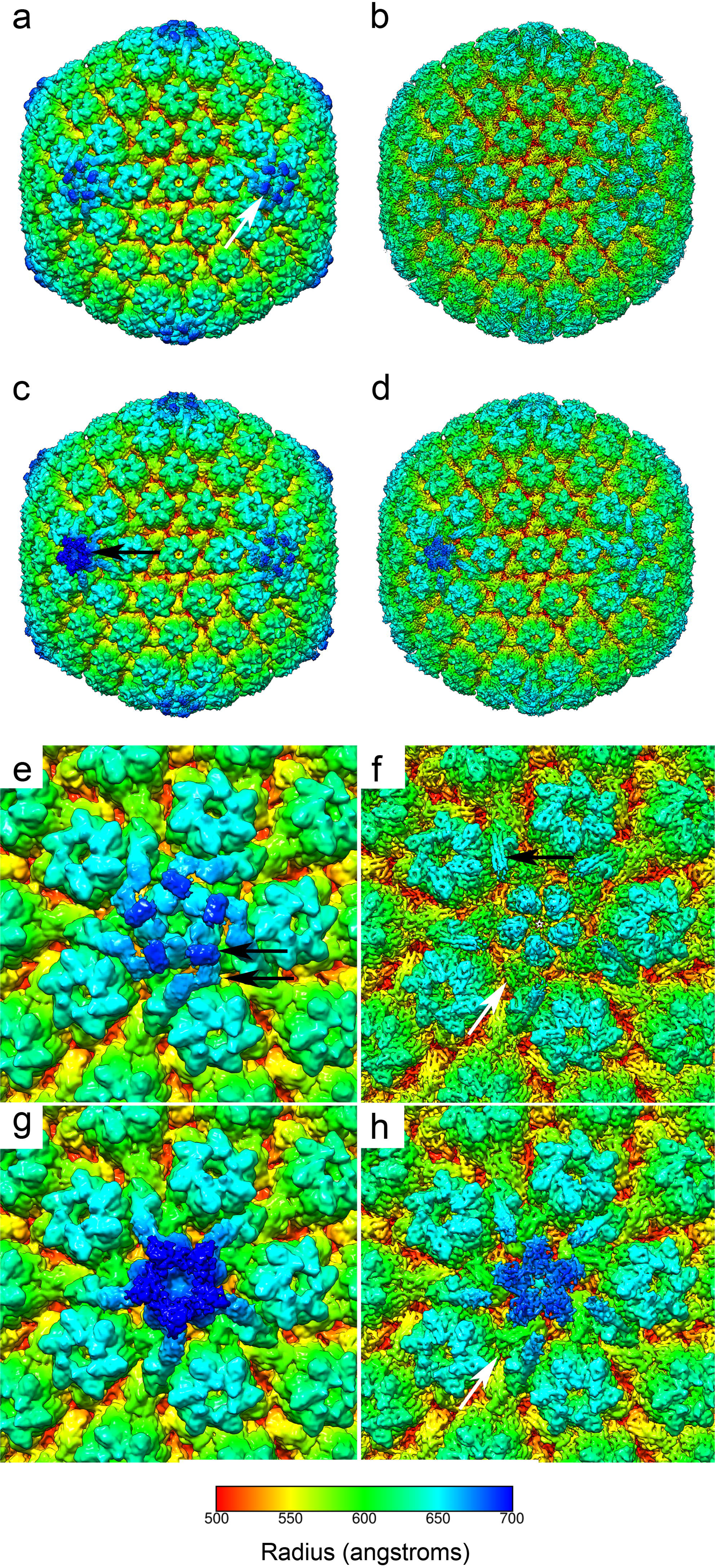
CryoEM and 3D image reconstruction of HSV-1 virions. Imposition of full icosahedral symmetry led to the calculation of a map at 6.3 angstroms resolution(a,b). At each five-fold symmetry axis the capsid associated tegument complex (CATC) binds to peri-pentonal triplexes to form an assembly that lays over the major-capsid protein penton (white arrow, a). A sharpened map reveals higher-resolution features and in particular the CATC four-helix bundle (b,f). Focussed classification led to the calculation of a C5 symmetric reconstruction at 7.7 angstroms resolution (c,d). This revealed the structure of the unique portal-vertex (c – black arrow). In the sharpened map, the CATC four-helix bundle is also well resolved at both portal and penton vertices (d,h). A close-up view of the penton-vertex highlights the structure of the CATC in particular two globular densities that have been attributed to the C-terminal domain of pUL25 (black arrows, e). In the sharpened map (f), the CATC four-helix bundle is highlighted (black arrow), as is the Ta triplex (white arrow). A close-up view of the portal-vertex shows that the five-fold symmetry axis is capped by a tiered structure comprising two star-shaped densities (g). The arms of the CATC are also angled more towards the five-fold axis. The sharpened map, viewed at a higher threshold does not show the distal tier of density (h). Interestingly, we see the position occupied by the Ta triplex at the penton vertex is occupied by a much larger globular density at the portal-vertex (white arrow). All maps are coloured according to radius in angstroms (see colour key).

Following the refinement of our icosahedral reconstruction, focussed classification was performed to identify and reconstruct the unique portal-vertex. A three-dimensional cylindrical mask was created to focus the 3D classification analysis onto a single five-fold symmetry axis. A metadata file was created with expanded symmetry such that each particle image had 60 orientations assigned, corresponding to the 60-fold redundancy of an icosahedral object. These data were subjected to 3D classification to calculate ten 3D reconstructions of the masked five-fold vertices, grouping the data into self-similar classes. A single class was identified that showed density significantly different from the known structures of penton-vertices. Interrogation of the metadata for this class showed that the majority of particle images contributed five views (figure S3). This is consistent with there being a single portal structure per virion, and as a consequence C5 symmetry is imposed on our reconstruction by the data.

3D reconstruction of this dataset led to the calculation of a map at a resolution of 7.7 Å (figure 1c-d, S4). This revealed a uniquely structured capsid-associated tegument assembly at the portal-vertex (figure 1g-h, movie S1), giving rise to the tail-like features previously described at lower resolution, termed the portal-vertex associated tegument (PVAT)^30^. At intermediate resolution we see that the portal-vertex CATC density closely resembles that seen at the penton-vertices. The four-helix bundle, comprised of pUL17, pUL25 and pUL36 is well-resolved, but is rotated approximately 15° counter-clockwise relative to that of the penton-vertex. In penton-vertices this part of the CATC is anchored to the capsid by binding to two triplexes, the peri-pentonal Ta triplex, and the Tc triplex. At the portal-vertex, the CATC still binds to the Tc triplex, however the Ta triplex is replaced with a globular density that appears almost twice the size of a normal triplex (white arrow, figure 1f,h). At this resolution, it is unclear whether this feature is made up of a heterotrimer of pUL38 and pUL18, plus an additional component, or whether the triplex has been entirely substituted by another protein. A likely candidate for this additional protein density is pUL36. This gene-product was recently shown to be a component of the CATC four-helix bundle, however only a small proportion of that large (3,164 amino-acids) protein has thus-far been accounted for^17^.

As well as the novel triplex-like density, we see major differences in the arrangement of the C-terminal domains of pUL25 at the portal-vertex. Peri-pentonal CATC complexes have two globular densities that bind to penton pUL19, one on top, and one to the side of each copy of the major capsid protein (black arrows, figure 1e). Portal-vertex CATC also shows ten globular densities that we attribute to the pUL25 C-terminal domains. These are however arranged to form two star-shaped tiers to make up the tail-like PVAT. The distal (outermost) tier being rotated ∼36° relative to the proximal one (movie S2). As was seen in our icosahedral reconstruction, the C-terminal domains of pUL25 are less well resolved than the CATC four-helix bundle; in particular the distal tier is only visible at lower threshold levels and is more clearly seen in unsharpened maps. This suggests that this feature is not rigidly constrained.

A central slice through the five-fold symmetric reconstruction of the herpes-simplex virion reveals the internal features of the portal-vertex and the packaged DNA (figure 2a-b, movie S3). Lying just inside the capsid shell we can see noisy density that we attribute to the portal protein pUL6 (Figure 2c-d). Single particle 3D reconstruction of this assembly has previously shown it to have cyclic symmetry forming oligomers ranging from undecamers to tetradecamers. Authentic pUL6 portals assemble as dodecamers^31^. Thus, a symmetry mismatch between this structure and the C5 capsid leads to incoherent averaging and explains the lack of clear density seen in our reconstruction. Nonetheless, we can segment this feature from our density map, to highlight its position and gross morphology (Figure 2e). Running through the centre of the portal-vertex, along the five-fold axis, we see strong density that extends through the portal and up against the inner surface of the PVAT. This is highlighted with a white arrow in figure 2a, and in movie S3. We interpret this as being the trailing end of the viral DNA, retained in the portal-vertex ready to be ejected upon infection of a new host cell. A similar feature has been shown in the bacteriophages Ø29 and Spp1, where specific interactions between stopper proteins and the DNA hold the end of the DNA within the tail assembly^32,33^.

**Figure 2.**
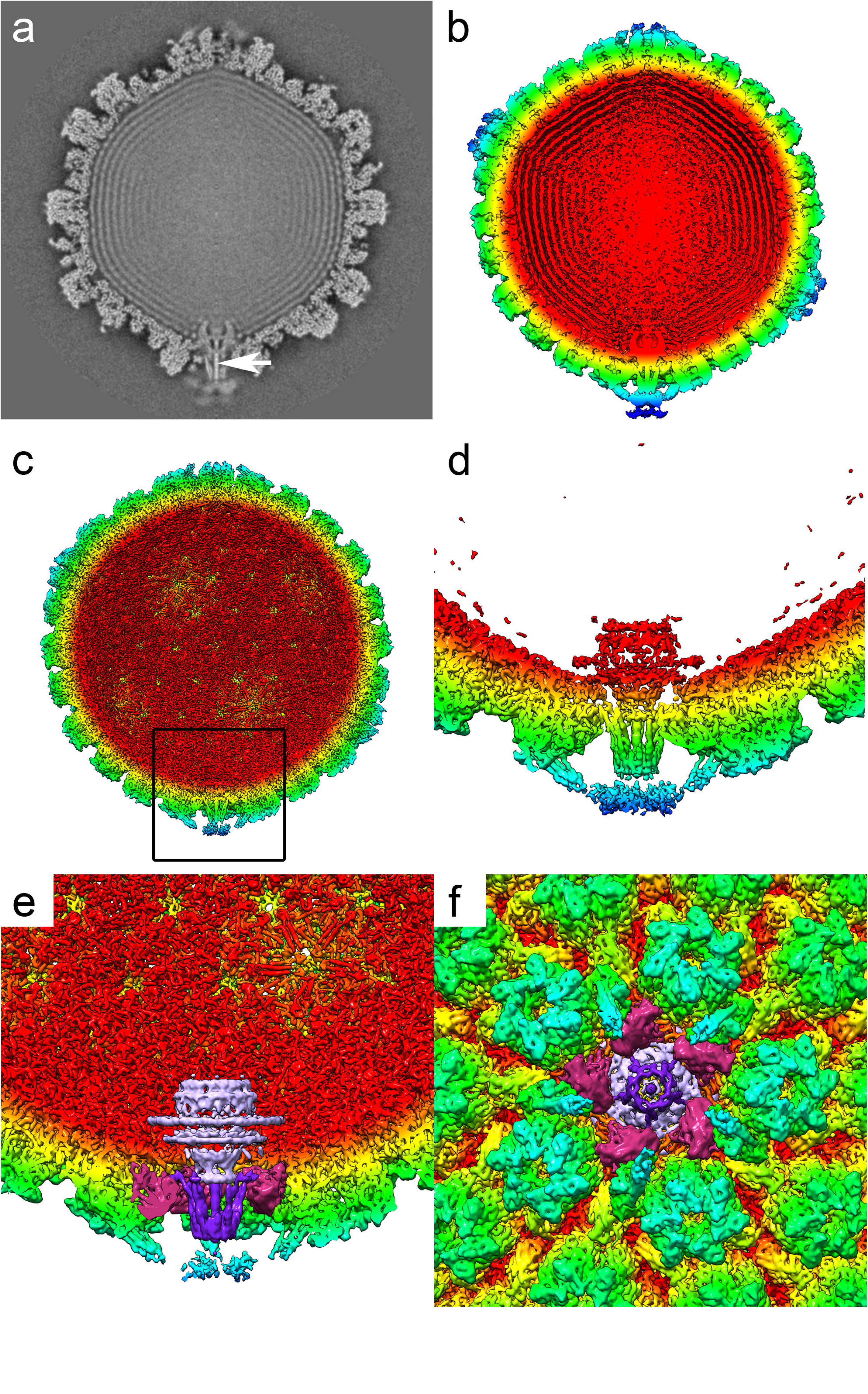
A central slice through the C5 reconstruction of the HSV-1 virion reveals the internal features of the portal-vertex (a). Notably a strong linear density is seen to run through the portal-vertex that we attribute to genomic DNA (white arrow). The outermost feature, the portal-vertex associated tegument (PVAT) is weakly resolved fuzzy density, suggesting that this feature is not well-constrained. Isosurface representation of the unsharpened map presents a clearer representation of the PVAT (b), while in the sharpened density map the packaged DNA is not seen, revealing the interior features of the capsid shell (c). A clipped, close-up view of the portal-vertex (boxed in c) highlights the morphology of the portal (pUL6), and lying between the portal and the PVAT, the pentameric portal-vertex protein, showing density consistent with two-helix coiled coil motifs (d). The density map was segmented to highlight three features, the portal (mauve), the pentameric portal-vertex protein (purple), and the peri-portal triplex-like density (magenta). The segmented portal-vertex is viewed both perpendicular to (e) and along (f) the five-fold symmetry axis.

Lying between the portal and the pUL25 PVAT density, a novel five-fold symmetric assembly replaces the usual pUL19 penton. This structure is composed of five protein subunits that are largely α-helical. In each subunit, a well resolved two helix bundle extends radially and is approximately 10nm in length (Figure 2e). This structure is anchored to the capsid through an interaction with the Ta triplex-like structures that are arranged about the portal-vertex. At this resolution, we see no evidence of an interaction with the major capsid protein. The identity of this portal-vertex protein is unclear. However, as the locations of all the components of the icosahedral capsid shell and the pUL6 portal protein are known, together with the CATC proteins, pUL17 and pUL25, a likely candidate is that this density is also the inner tegument protein, pUL36. As noted above pUL36 is known to be a CATC component, however this represents only a small proportion of this large protein, leaving the rest unaccounted for.

Herpesviruses are structurally and biologically similar to the caudovirales – DNA containing tailed bacteriophages. It has been suggested that these two viral groups share a common ancestry, owing to fold-conservation in the major capsid protein (pUL19 has a fold widely seen in tailed phage capsid proteins and first described in HK97)^34^, sequence similarities involving the gene encoding pUL15 terminase protein^35^, and their similar capsid assembly and DNA packaging strategies. Like herpesviruses, tailed phages assemble empty procapsids. The viral genome is pumped into the procapsid through a unique portal-vertex containing a dodecameric motor protein leading to capsid maturation. In tailed phages this process is mediated by terminase complexes that usually comprise two proteins. A small subunit, mediates binding of the terminase to the viral DNA. A large subunit, having both endonuclease and ATPase activity powers the translocation of the genome into the capsid through the portal, and then cleaves the DNA when the head is full, or a cleavage signal is detected. The HSV-1 terminase is made up of three proteins, pUL15, pUL28 and pUL33^15^. pUL28 is known to bind viral DNA, playing the role of the small terminase subunit^36^. pUL15 shows sequence similarity to the large terminase subunit of phage and is predicted to have both endonuclease and ATPase activity. The function of pUL33 is rather less well defined. It is predicted to be largely α-helical in structure^37^ and thus we do not rule out the possibility that the unidentified portal-vertex protein could be pUL33. By analogy to bacteriophage terminase complexes however, the terminase assembly is expected to detach from the portal following cleavage of the concatamer, remaining associated with the unpackaged DNA, ready to engage the portal in another empty capsid. Indeed, proteomics analysis has failed to detect terminase components in purified HSV-1 virions. Further biochemical and structural analysis of isolated nucleocapsids and virions is therefore required to unambiguously identify this protein.

In addition to the much-improved view of the HSV-1 portal-vertex, our data also reveal detailed structural information for the packaged DNA. Our low-symmetry reconstruction reveals that the DNA is packed as a left-handed spool (figure 3, movie S4). The outer-most three layers of DNA clearly show an identical orientation of the spooled DNA. The presence of clear density for the genomic DNA indicates that a very consistent packaging process occurs when DNA is pumped into the nascent nucleocapsid. Our data suggest that the bending forces on the incoming DNA, combined with lateral or rotational forces imparted by the portal motor, lead to a consistent orientation for the DNA spool inside every virion. Herpesvirus DNA is packed to a very high density^38^, indeed in another herpesvirus, cytomegalovirus, genome density approaches the limits of what may occur without transition to a crystalline state^39^. It is known that both pUL25 and pUL36 are critical to the retention of packaged DNA. Our data suggest a reason for this, showing that pUL25 forms a double-layered cap on the outer-face of the portal-vertex (the PVAT). We can see strong density, that we interpret as being DNA, running through the central channel of the portal-vertex, extending through both the pUL6 portal and the unidentified (possibly pUL36) pentameric portal-vertex protein complex, reaching to the distal tip of the latter structure, and terminating at the position occupied by the PVAT/pUL25 density.

**Figure 3.**
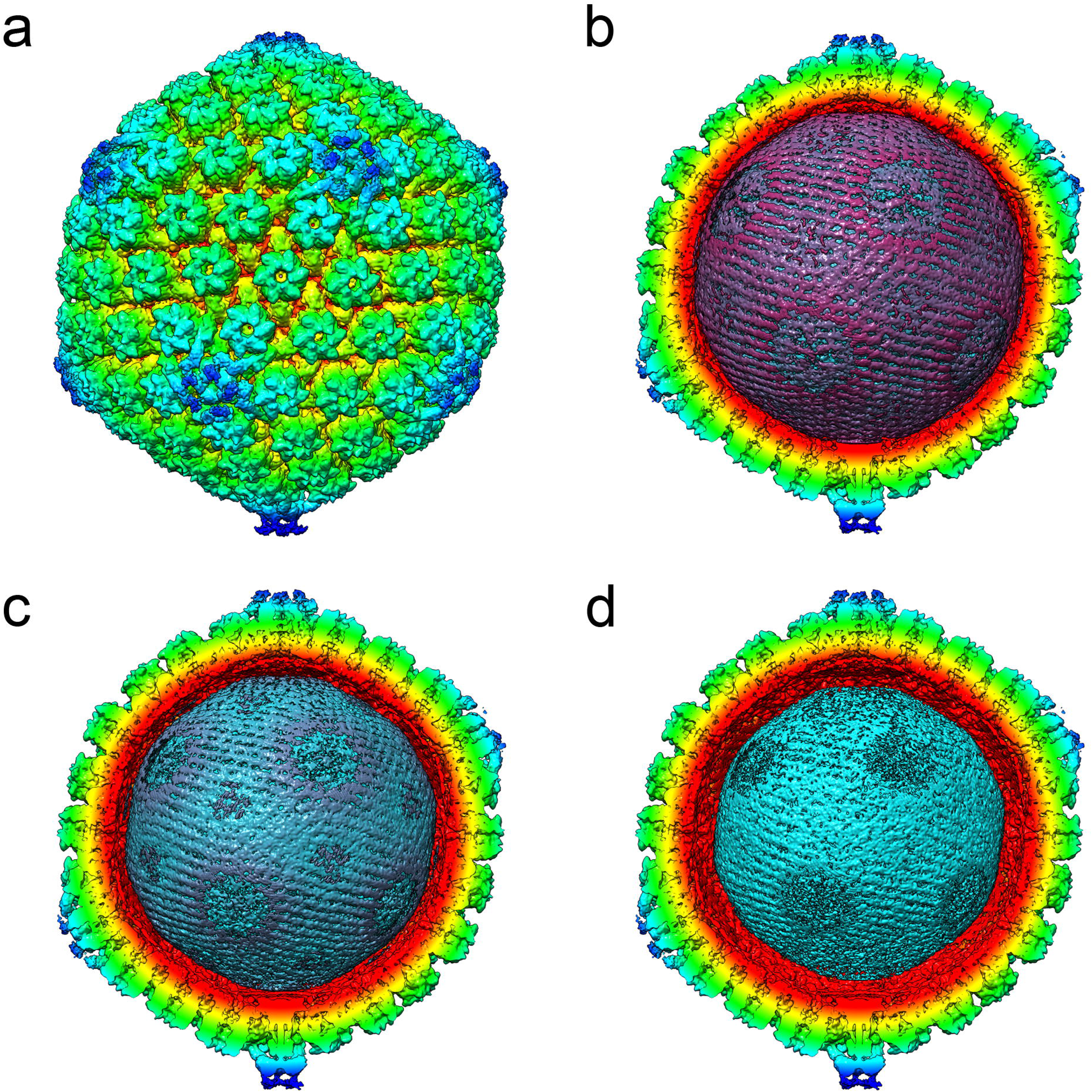
The unsharpened C5 reconstruction of HSV-1 showing the entire nucleocapsid (a) and a series of radially cropped views of the interior density, reveal the highly-ordered arrangement of packaged DNA. The outermost (b), second (c) and third (d) shells are shown, revealing a left-handed spool of density.

## Summary

We have applied a novel image processing approach to determining the structure of an important asymmetric feature of the icosahedral herpesvirus nucleocapsid. Our data provide new insights into the organisation and composition of this important molecular machine. The herpesvirus portal-vertex is the unique site through which the viral genome both enters and exits the nucleocapsid. It is therefore a critical assembly in both virion morphogenesis and initiation of infection. We show that the portal-vertex associated tegument, a tail-like structure previously described at low resolution, is likely composed of the C-terminal domain of pUL25, a protein known to be critical to DNA retention. We have also described two novel structures. The first, a hitherto unknown pentameric assembly, lays between the portal and PVAT in place of the pUL19 penton and has a clear two-helix coiled-coil structure. The second is a large assembly that occupies the space usually taken up by the Ta triplex in peri-pentonal vertices. We have suggested possible identities for both structures, however further biochemical analysis, coupled to higher resolution structure determination are required to provide a definitive description. In agreement with previous lower-resolution studies that employed electron tomography, we show that the portal motor, comprised of twelve copies of pUL6, lays against the inner edge of the capsid shell. Finally, our data show that HSV-1 genomic DNA is packaged as a left-handed spool, arranged in concentric layers. This suggests that the process of genome translocation combined with interactions between the inner surface of the capsid and successive layers of DNA lead to a highly reproducible packing process. This is perhaps not as remarkable as one might initially surmise. Herpesviruses package their genomes to extremely high density. It is critical then, that the genome should be able to reliably achieve this high density, and with equal reliability eject the complete genome through the nuclear-pore to initiate infection.

## Materials and Methods

### Culture and purification of HSV-1 virions

Herpes simplex virus was propagated in BHK cells grown in roller bottles containing growth medium (GMEM, 10%FCS, 10% tryptose phosphate broth). Confluent cells were infected with HSV-1 strain 17syn+ at a m.o.i. of 0.002 pfu/cell. Cells were incubated at 37°C for 3 days then harvested by shaking into the medium. Cells and media were centrifuged at 1600×g for 10 minutes to remove cell debris. The supernatant was then transferred to a new tube and spun at 17,000×g for 2 hours. The pellet was then gently resuspended on ice overnight by overlaying with 2ml GMEM. The resuspended material was removed to a new tube and clarified by spinning at 200×g for 10 minutes. The supernatant was layered onto the top of a 5-15% Ficoll gradient prepared in GMEM and centrifuged at 26,000×g for 2 hours. The opaque band containing the virions was collected by side puncture of the tube using an 18-gauge needle. Collected virions were diluted in GMEM and pelleted at 40,000×g for 1 hour. The pellet was washed gently in PBS and then allowed to resuspend in 50-100ul PBS by incubating on ice for at least 1 hour.

### Cryo-electron microscopy

HSV virions were prepared for cryo-EM by plunge freezing into liquid ethane. 4 μl of purified virions was loaded onto a freshly glow-discharged holey-carbon support film (R2/2 Quantifoil), in an FEI Vitrobot mk. IV vitrification robot. The grid was immediately blotted for 3 seconds and then plunged into liquid nitrogen cooled liquid ethane. Grids were imaged at the UK national cryoEM facility (electron bioimaging centre – eBIC) at Diamond Light Source, Harwell, in an FEI Titan Krios cryo-transmission electron microscope at a nominal magnification of 81,000×. Images were recorded as ‘movies’ on a Falcon III camera, operated in integrating mode with sampling of 1.78 Å/pixel. Movies were recorded as 12 second exposures giving a total of 480 detector frames that were integrated into 40 fractions, with a dose rate of 1.95 e/Å/fraction.

### Image processing

All image processing was performed using Relion 2.1^40^, on a GPU workstation running Linux CentOS 7. 3,702 micrograph movies were processed to correct for particle movement using Motioncor2^41^, and the defocus for each motion-corrected micrograph was estimated using GCTF^42^. A small subset of particles was picked and subjected to 2D classification, to produce a template for automated particle picking, which yielded a total of 12,431 virion images for further processing. Particles were initially extracted with 2× binning. After 2D classification of this dataset 7,476 particles were selected for *ab initio* calculation of a starting model followed by 3D classification, imposing full icosahedral symmetry in both cases. This led to the definition of a final dataset of 6,069 virion images that were taken forward for 3D refinement with full icosahedral symmetry. Once completed, a new dataset of particles with 1.5× binning was extracted and used to calculate the final icosahedral reconstruction.

Following icosahedral reconstruction, focused classification was performed to identify the unique portal-vertices^43^. A cylindrical mask was prepared in SPIDER^44^, to cover a single five-fold symmetry axis. A metadata file (STAR file) was generated to expand the symmetry for our dataset, i.e. for each virion image, 60 orientations were defined corresponding to the 60 symmetry related views of the icosahedral object. To speed up the calculation a new dataset was extracted from the raw micrographs with 5× binning. These data were then subjected to masked classification, to reconstruct a single five-fold vertex and classify the data into self-similar classes. During this process orientations and origins were not refined. A total of 10 classes were calculated, one of which was identified as containing the unique portal-vertex. To calculate our final C5 symmetric reconstruction, we used 1.5× binned particle images.

Resolution assessment was performed in Relion, using the postprocessing task to mask the density maps and calculate the ‘gold-standard’ Fourier shell correlation. A B-factor was estimated and applied to each reconstruction^45^, which was then interpreted by visualization in UCSF Chimera^46^.

The C5 reconstruction has been deposited in the EM databank with accession code EMD-4347.

## Acknowledgements

The authors acknowledge Diamond Light Source for access and support of the Cryo-EM facilities at the UK national electron bio-imaging centre (eBIC), proposal EM16637-4, funded by Wellcome, the Medical Research Council and the Biotechnology and Biological Sciences Research Council. In particular we thank Dr Kyle Dent for his expert support in collecting the data described herein. This work was supported by the Medical Research Council (MC_UU_12014/7).

