## Supplementary Materials for "Structure of the herpes-simplex virus portal-vertex"

### Short communication

Marion McElwee, Swetha Vijayakrishnan, Frazer Rixon and David Bhella

#### Supplemental data

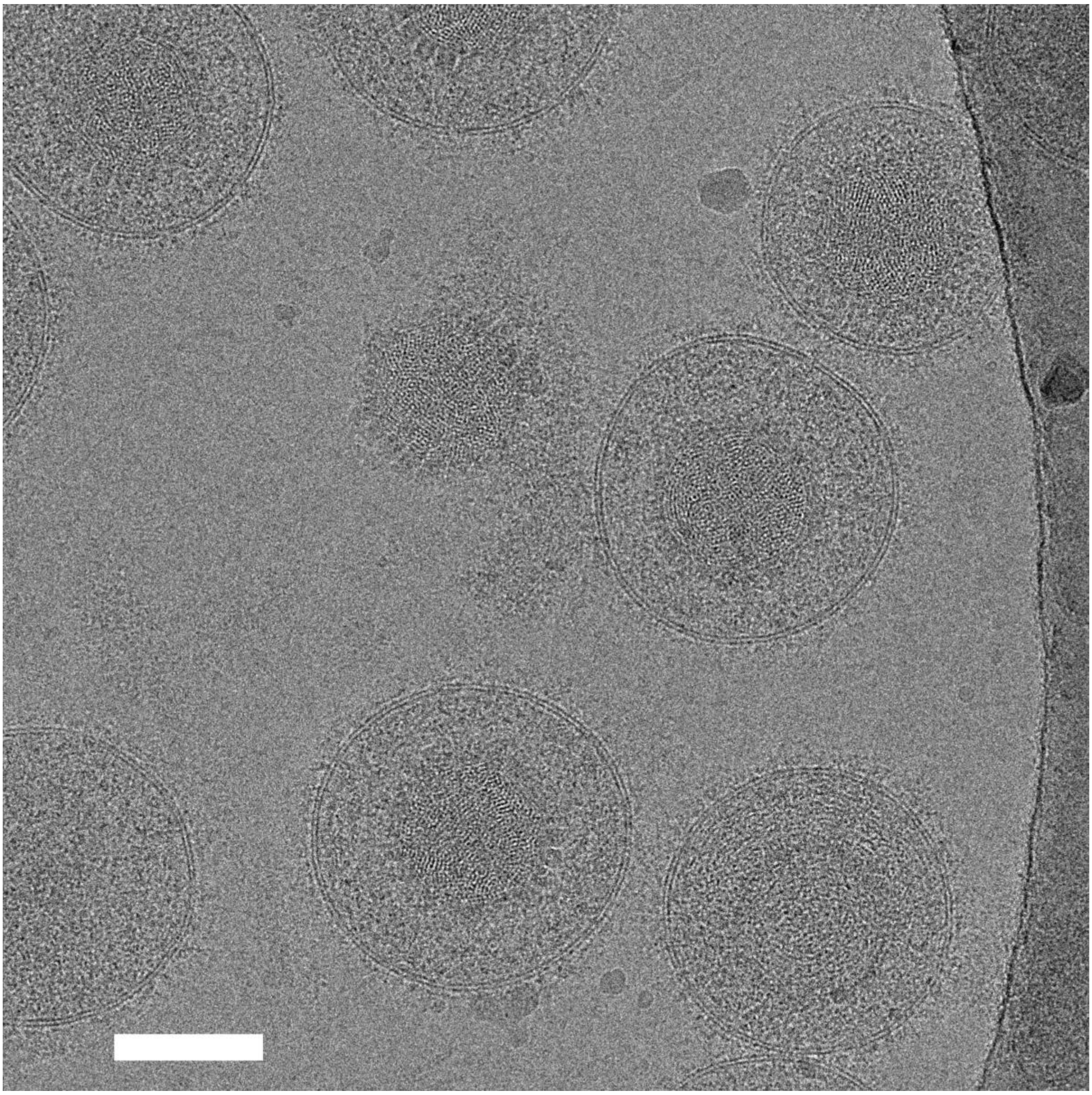

**Figure S1**
Cryomicrograph of HSV-1 virions. Scale bar = 100nm

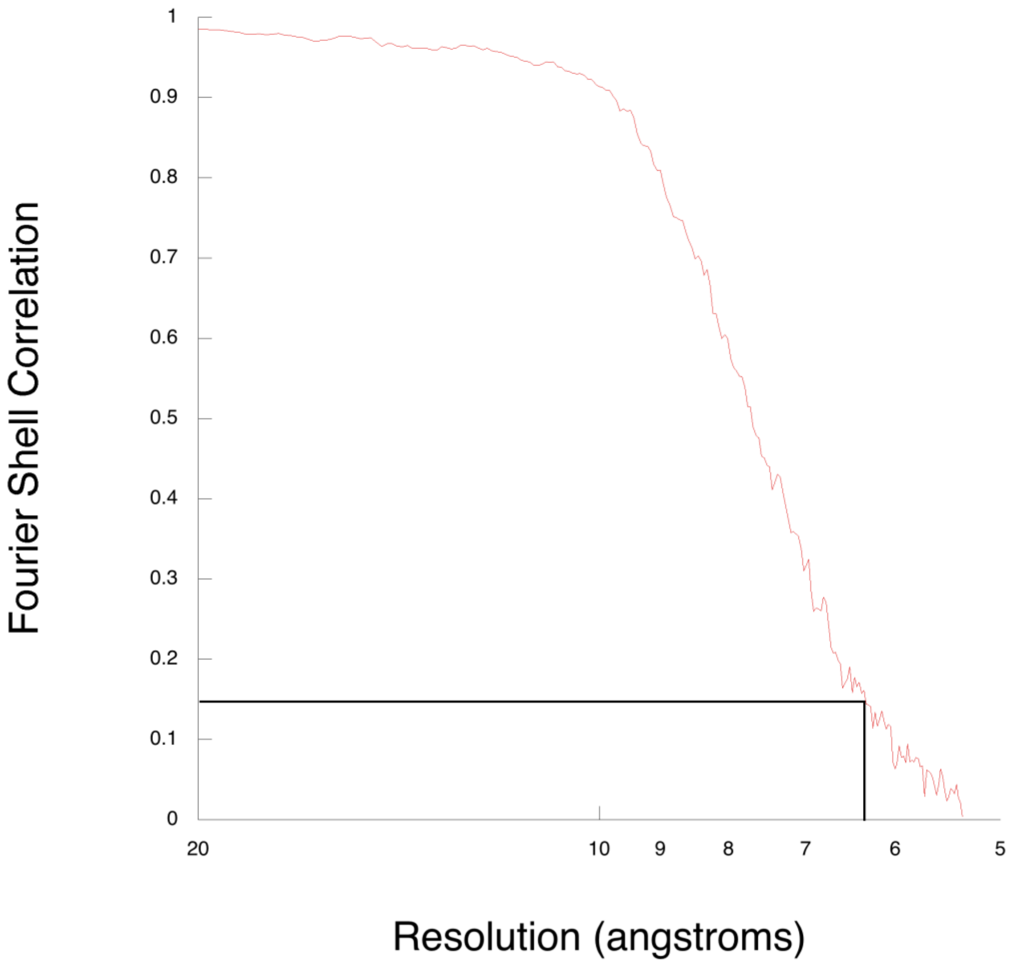


**Figure S2**

*Fourier shell correlation plot for the icosahedral reconstruction of HSV-1*

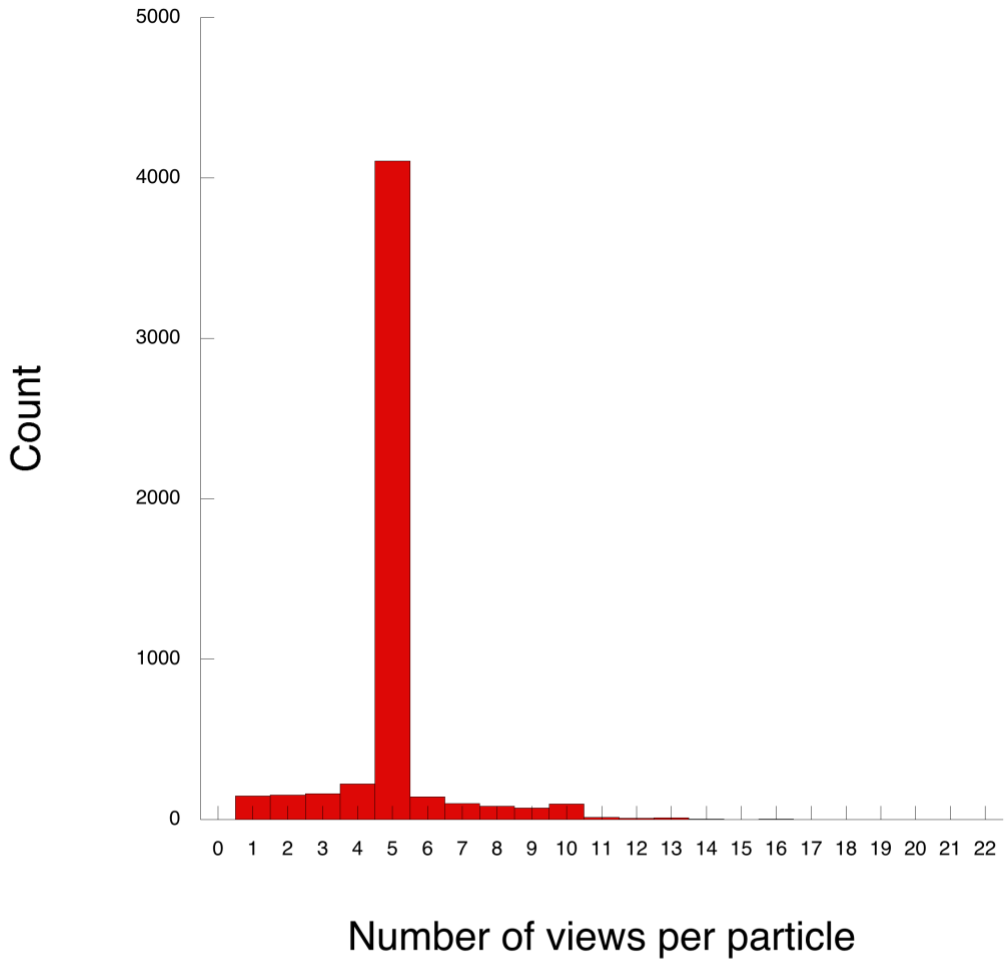


**Figure S3**

*Focussed classification to identify the portal-vertex in HSV-1.* To solve the structure of the portal vertex, focussed classification was used. This allows us to determine the structures of asymmetric features in high-symmetry objects. To achieve this, we expand the symmetry of the dataset such that each particle image has multiple orientations specified according to the redundancy of the symmetry group. In the case of icosahedral objects, each particle would contribute 60 views. A meta-data file was therefore created in which each particle image was assigned 60 symmetry-related orientations based on the single orientation that had been determined during 3D reconstruction with full icosahedral-symmetry. Masked 3D classification, focussing on a single five-fold axis led to the definition of a class that showed density significantly different from the known penton-vertex structure. To understand the distribution of particle views present in this class, we sought to determine the number of times each particle was assigned to it. From our dataset of 6,069 virion images we produce a metadata file containing 364,140 putative views (60×6,069). The portal-vertex class was found to contain 26,891 entries, ~7.4% of the total dataset. There are twelve five-fold symmetry axes in an icosahedral object, thus we would expect 8.3% (1/12) of the data to be assigned to the unique portal vertex. Further interrogation of the metadata file for this class revealed that 5,337 unique particles were represented in the data-set, thus 732 particles did not present portal density that was readily identified by our analysis. We determined the number of views for each virion image in the portal vertex class, this revealed that the median number of views per particle was 5, consistent with the C5 symmetry of the portal axis and indicating that most (if not all) particles have only one portal.

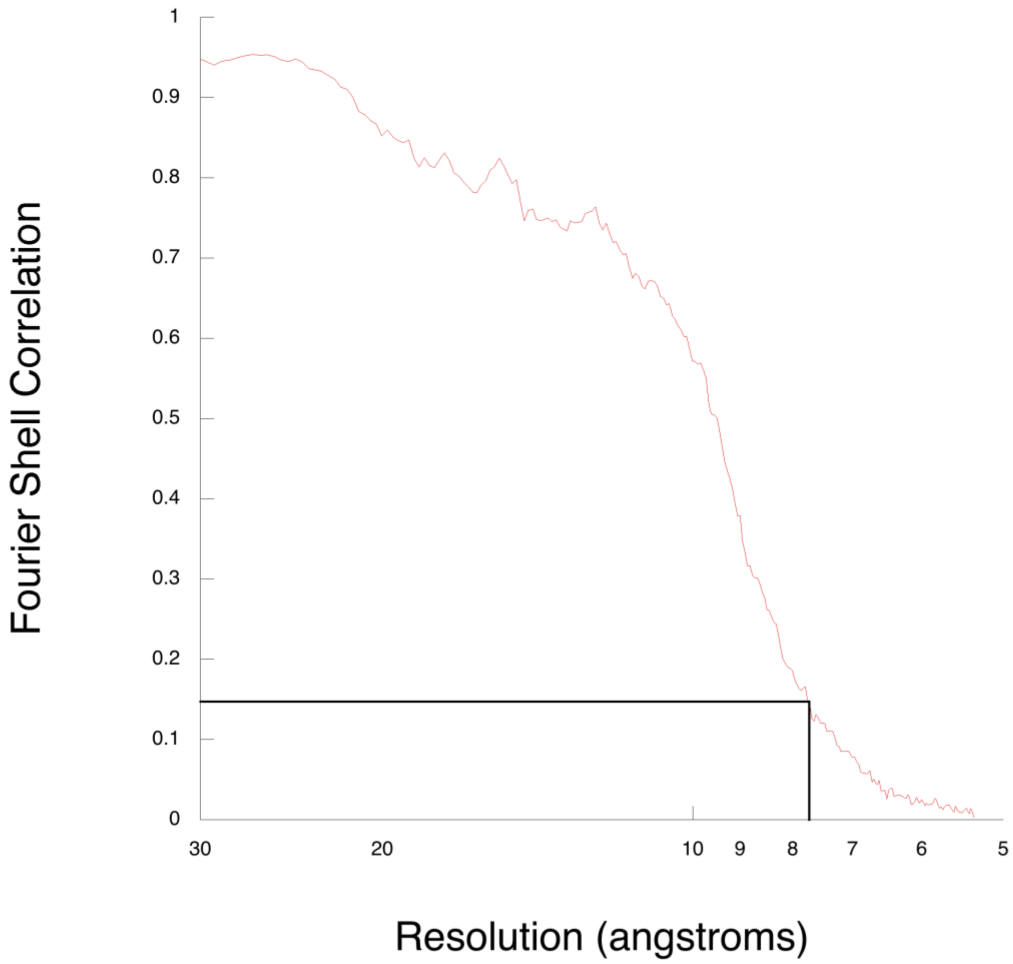


**Figure S4**

*Fourier shell correlation plot for the C5 reconstruction of HSV-1*
